# Stags, Hawks, and Doves: Social Evolution Theory and Individual Variation in Cooperation

**DOI:** 10.1101/126367

**Authors:** Jeremy Van Cleve

## Abstract

One of the triumphs of evolutionary biology is the discovery of robust mechanisms that promote the evolution of cooperative behaviors even when those behaviors reduce the fertility or survival of cooperators. Though these mechanisms, kin selection, reciprocity, and nonlinear payoffs to cooperation, have been extensively studied separately, investigating their joint effect on the evolution of cooperation has been more difficult. Moreover, how these mechanisms shape variation in cooperation is not well known. Such variation is crucial for understanding the evolution of behavioral syndromes and animal personality. Here, I use the tools of kin selection theory and evolutionary game theory to build a framework that integrates these mechanisms for pairwise social interactions. Using relatedness as a measure of the strength of kin selection, responsiveness as a measure of reciprocity, and synergy as a measure of payoff nonlinearity, I show how different combinations of these three parameters produce directional selection for or against cooperation or variation in levels of cooperation via balancing or diversifying selection. Moreover, each of these outcomes maps uniquely to one of four classic games from evolutionary game theory, which means that modulating relatedness, responsiveness, and synergy effectively transforms the payoff matrix from one the evolutionary game to another. Assuming that cooperation exacts a fertility cost on cooperators and provides a fertility benefit to social partners, a prisoner’s dilemma game and directional selection against cooperation occur when relatedness and responsiveness are low and synergy is not too positive. Enough positive synergy in these conditions generates a stag-hunt game and diversifying selection. High levels of relatedness or responsiveness turn cooperation from a fitness cost into a fitness benefit, which produces a mutualism game and directional selection for cooperation when synergy is not too negative. Sufficiently negative synergy in this case creates a hawk-dove game and balancing selection for cooperation. I extend the results with relatedness and synergy to larger social groups and show that how group size changes the effect of relatedness and synergy on selection for cooperation depends on how the per capita benefit of cooperation changes with group size. Together, these results provide a general framework with which to generate comparative predictions that can be tested using quantitative genetic techniques and experimental techniques that manipulate investment in cooperation. These predictions will help us understand both interspecific variation in cooperation as well as within-population and within-group variation in cooperation related to behavioral syndromes.

## 1. Introduction

Evolutionary biologists have long been fascinated by the ability of natural selection to craft complex and often highly cooperative social behaviors. These behaviors are often costly to perform and include, among many other examples, the production of fruiting bodies by the slime mold *Dictyostelium discoideum* whose stalks sacrifice themselves for the survival of the spores (Strassmann et al., 2000), the formation of termite mounds with millions of individual (Korb, 2010), and the formation of human towers up to nine levels high as cultural symbols in Catalonia (Vaczi, 2016). While high levels of cooperation have often been described as an evolutionary “difficulty” (Darwin, 1859) or “puzzle” (Colman, 2006) when such cooperation exacts fitness costs, multiple biological avenues exist that lead to the evolution of significant levels of cooperation. These avenues include kin selection, reciprocity, and nonlinear payoffs for investing in cooperation.

Hamilton pioneered the exploration of kin selection (Hamilton, 1964, 1970) and showed how cooperative behaviors can experience positive selection if the individuals who receive the benefits share genes with the individuals who generate the benefits. In the years since Hamilton’s original insight, kin selection has developed into a robust theoretical approach (Frank, 1998) grounded in population genetics (Rousset and Billiard, 2000; Rousset, 2004; Van Cleve, 2015) and applicable to species with complex life histories (Taylor, 1992; Taylor and Irwin, 2000; Lehmann and Rousset, 2010, 2014; Van Cleve, 2015) and population structures (Rousset and Billiard, 2000; Lehmann et al., 2007; Taylor et al., 2007; Ohtsuki and Nowak, 2008; Tarnita et al., 2009; Taylor, 2016). Another important avenue is the repeated and reciprocal exchange of help between individuals, which is often called reciprocity (Trivers, 1971; Axelrod and Hamilton, 1981) or responsiveness (Akçay et al., 2009; Akçay and Van Cleve, 2012; André, 2015). Though relatedness and responsiveness operate on much different timescales (evolutionary and behavioral, respectively), they both generate positive selection for cooperation by correlating among individuals the benefits and costs of cooperation (Queller, 1985). Finally, the payoffs of cooperation can shift in a nonlinear way due to changes in the underlying ecology of the social interaction. For example, a troupe of chimpanzees might shift from foraging for fruit to hunting red colobus monkeys. This shift involves a nonlinear increase in the benefit of cooperating because hunting as a troupe can generate more food per chimpanzee than hunting alone. The additional nonlinear benefit in this example is sometimes called (positive) synergy (Queller, 1985; Frank, 1995; Hauert et al., 2006; Van Cleve and Akçay, 2014), and if its large enough, it can create selection for cooperation even when cooperation is individually costly.

Typically, each of the above avenues has been studied independently with a focus on how they affect evolved levels of cooperation. However, these avenues interact in important ways to change not only the evolved level of cooperation but also whether variation in levels of cooperation evolves and whether that variation is within or among groups. Within and among group variation in cooperation can be characterized as different behavioral syndromes or personalities (Sih et al., 2004; Wolf and Weissing, 2012; Jandt et al., 2014). In this paper, I will illustrate how the intersection of relatedness, responsiveness, and synergy can be studied systematically using the tools of kin selection and evolutionary game theory. Specifically, I will show how the scenarios of directional selection for or against cooperation and selection for variation in cooperation within or among groups correspond to the fitness payoffs of different social games from evolutionary game theory. Changing the fitness payoffs and shifting from one game to another occurs by changing the level of relatedness, responsiveness, or synergy. I show that the strongest effect of relatedness and responsiveness is turning directional selection against cooperation to direction selection for cooperation. Synergy is most important for determining whether variation in cooperation evolves and whether it is among or within groups: positive synergy is generally required for variation among groups and negative synergy for variation within groups. Finally, I show increasing the size of the social group can have a strong effect on creating selection for or against variation in cooperation depending on whether the per capita benefit of cooperation shrinks with group size or not.

## 2 Types of natural selection and evolutionary games

Regardless of how natural selection on investment in cooperation is generated, it can be classified as either directional, diversifying, or balancing, and each of these kinds of selection corresponds to a two-player game from evolutionary game theory (Cressman, 2003). Suppose that two individuals engage in a social interaction (which is known as a “game” in game theory) where either individual can invest in cooperation (C) or not invest in cooperation (D). The payoffs for each combination of actions is given by the payoff matrix in Table 1. Cooperation exacts an individual cost *C* from the cooperating individual and produces an additive benefit *B* available to a social partner. Mutual cooperation produces an additional payoff D, which is the payoff synergy. For the purposes of studying a cooperative or helping behavior, I assume that the benefit of cooperation is positive, *B* > 0, and that the benefit is larger than the cost, *B* > *C*.

**Table 1:** Payoff matrix for a pairwise or two-player game.

|  | C | D |
| --- | --- | --- |
| Payoff to C | $B - C + D$ | $-C$ |
| Payoff to D | $B$ | $0$ |

It is important to specify the units of the payoffs in Table 1 since these units determine what parts of the life cycle are and are not packed into these payoffs. The units are always direct or proximate measures of a component of fitness, namely survival or fertility. The life cycle of an organism includes mating and reproduction and can include dispersal, density-dependent regulation, and social interactions at any stage between newborn and adult stages. For example, if the payoff unit is a proxy for how the social interaction affects the survival of juveniles to the adult stage, then predicting what kind of selection there is on cooperation will also depend on information about survival rates of adults, fertility, and any potential population structure generated by limited dispersal. In order to start with the simplest possible biological example, I assume that the units of payoff are proxies for fertility whereas survival is constant and unaffected by the social interaction. Additionally, individuals are asexual, live in a single population of large size, reproduce once, and are immediately replaced by their offspring (semelparity). Given this simple scenario, the payoffs in Table 1 can fully characterize the type of selection on cooperation.

Positive directional selection corresponds to a mutualism game (Clements and Stephens, 1995) where individuals maximize their payoff by cooperating no matter what their partners do. This occurs when there is an individual benefit to cooperation (i.e., the cost is negative, *C* < 0) and synergy is positive, zero, or not too negative (*D* > *C*). Positive directional selection for cooperation leads to all individuals cooperating and no variation in the level of cooperation. For example, predator avoidance behaviors can be mutualistic since avoiding a predator has an individual benefit and can provide useful information to social partners about the presence of a predator.

Negative directional selection leads to no individuals cooperating and produces no variation in the level of cooperation. The prisoner’s dilemma game (Rapoport and Chammah, 1965) generates negative directional selection for cooperation and occurs when cooperation has an individual cost (*C* > 0) and synergy is negative, zero, or not too positive (*D* < *C*). An example of a prisoner’s dilemma is the rate of consumption of a finite resource such as ripe fruit from a tree located by a group of chimpanzees (Chapman et al., 1995). Eating the fruit faster yields a short-term benefit for each individual but results in the fruit running out more quickly when all individuals eat quickly; eating the fruit more slowly has a short term cost for each individual but benefits the group by maintaining the resource further into the future.

Diversifying selection is produced by a stag-hunt (or coordination) game, which occurs when an individual obtains a higher payoff by choosing the same action, cooperate or do not cooperate, as its partner (Skyrms, 2001). This occurs when cooperation is individually costly (*C* > 0) and synergy is sufficiently positive (*D* > *C*). The stag-hunt game has two solutions (Nash equilibria in game theory terminology), both individuals cooperate and neither individual cooperates, and which outcome arises depends on the organism’s decision making process. Simple decision making processes that attempt to increase an organism’s payoff lead all individuals to cooperate in some populations and no individuals to cooperate in other populations, and this generates among group variation in cooperation. Chimpanzees hunting red colobus monkeys could correspond to a stag-hunt game since larger groups, which have more investment in cooperation, are more successful than smaller groups (Boesch and Boesch, 1989) that might more easily break up in favor of individuals hunting or foraging alone.

Finally, balancing selection occurs when individuals obtain higher payoffs when choosing the opposite action (cooperate or do not cooperate) of their partner. These payoffs yield a snowdrfit (Sugden, 2005) or hawk-dove game (Maynard Smith and Price, 1973; Maynard Smith and Parker, 1976) where cooperation is individually beneficial (*C* < 0) but obtaining payoff via mutual cooperation is inefficient compared to not cooperating and obtaining payoff for a cooperating partner (*B* – *C* + *D* < *B*). The latter condition implies negative synergy (*D* < *C* < 0). In the simplest scenario where individuals are randomly paired with social partners in a very large population, natural selection leads to all individuals cooperating randomly with probability 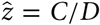 (called a mixed strategy Nash equilibrium in game theory). If individuals do not pick strategies randomly, a polymorphism between cooperating individuals and non-cooperating individuals can evolve where the frequency of cooperators is the mixed strategy probability of cooperation 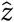 (Bergstrom and Godfrey-Smith, 1998). In either case, mixed strategy or polymorphism, each population has a mixture of cooperation and noncooperation and variation in cooperation is within groups. A hawk-dove game might describe how chimpanzees forage for fruit. Foraging is individually beneficial when no one else is doing it but locating fruit might also reveal that location to social partners who then receive the benefit without paying the cost of foraging (analogous to a producer-scrounger game; Barnard and Sibly, 1981; Vickery et al., 1991).

In order to see how the payoffs of the game determine the type of selection, suppose that a focal individual who cooperates with probability *z_•_* interacts with an individual who cooperates with probability *z*_∘_. Using Table 1, the payoff *π* obtained by the focal individual is given by

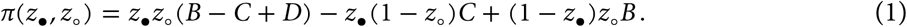

Further, assume that there are two alleles for cooperation in the population: a resident (wild type) allele that produces cooperation with probability *z* and a mutant allele that produces cooperation with probability *z* + *δ*. If *δ* is small (terms smaller than *δ* can be ignored), then the change in the frequency of the mutant allele, Δ*p*, is proportional to (denoted by the symbol “∝”) the change in the payoff the focal individual gets by increasing its probability of cooperating (Taylor and Jonker, 1978; Hofbauer and Sigmund, 1998),

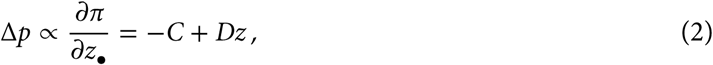
evaluated at the resident probability of cooperating, *z*. Equation (2) shows how the payoffs from the game and the probability of cooperating determine the kind of selection on the mutant: directional selection for increasing the frequency of the mutant (Δ*p* > 0) occurs when *–C* > 0 and *D* > *C*, directional selection for decreasing the frequency of the mutant (Δ*p* < 0) occurs when *C* > 0 and *D* < *C*, diversifying selection Δ*p* > 0 for *z* = 1 and Δ*p* < 0 for *z* = 0) occurs when *D* > *C* > 0, and balancing selection (Δ*p* > 0 for 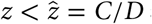 and 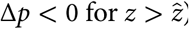) occurs when *D* < *C* < 0.

Table 1 summarizes how the different types of selection correspond to the four games and what level of cooperation and what kind variation in cooperation level each game supports. So far, I have shown how one can shift between these games by changing how synergy compares to the cost of cooperation. However, shifting between games also can occur by changes in relatedness and responsiveness. In order to understand how these three mechanisms can interact together to shift the fitness payoffs from one game to another, I use mathematical tools from kin selection and evolutionary game theory.

## 3 The direct fitness method and kin selection

The direct fitness method of kin selection theory (Frank, 1998; Rousset and Billiard, 2000) uses population genetics to derive an expression for the expected change in the frequency of a mutant allele, Δ*p*, when the population is spatially structured and local dispersal builds up genetic relatedness between neighboring individuals. The direct fitness method can account for arbitrarily complex populations, but to keep things simple I assume that individuals live in groups of equal size that are connected by some rate of dispersal. Moreover, individuals engage in a pairwise social interaction or game whose payoffs in units of fertility are given in Table 1. Assuming that the change in investment is small, one can show that (Hamilton, 1964, 1970; Taylor and Frank, 1996; Frank, 1998; Rousset and Billiard, 2000; Rousset, 2004)

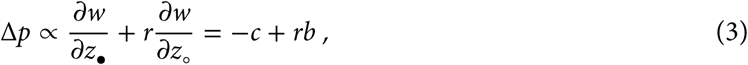
where *w* is the expected fitness of a focal individual, *r* is the genetic relatedness, *z*_•_ is the strategy of a focal individual, *z_∘_* is the mean strategy of other individuals in focal group, and the derivatives are evaluated at the resident strategy *z*. Equation (3) is Hamilton’s rule (Hamilton, 1964, 1970) and shows how the direct effect of selection, the fitness cost –*c*, and the indirect effect of selection, the fitness benefit *b* weighted by relatedness *r*, combine to determine the direction of selection on a social trait. The fitness cost is the change in expected fitness due to the effect of a change in the focal individual’s behavior, 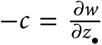, and the fitness benefit is the change in expected fitness due to a change in the strategy of other individuals in the social group, 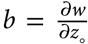. The expected fitness *w* measures the expected number of surviving offspring of a focal individual over a single generation and thus must account for how demographic factors such as dispersal and density dependent regulation affect fitness.

While Hamilton’s rule in Equation (3) is useful conceptually for partitioning the effects of selection on a social trait based on fitness and relatedness, it is less useful for showing how relatedness can shift a social interaction between the different game types in Table 2 as function of the payoffs in Table 1. This is because the payoffs are packaged within the fitness cost and benefit, *–c* and *b*, along with other demographic factors. A more useful expression of Δ*p* unpacks the payoffs from within *–c* and *b* and gathers the effects of demographic forces and relatedness into a single term. Using the assumptions necessary for deriving Equation (3), this expression is

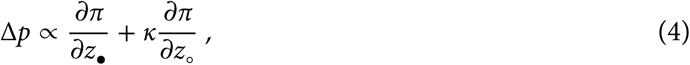
where *π* is the expected payoff the focal individual receives, the derivatives are evaluated at the resident strategy *z*, and *κ* is the “scaled-relatedness” coefficient (Lehmann and Rousset, 2010; Van Cleve and Lehmann, 2013; Van Cleve, 2015; Peña et al., 2015). The scaled relatedness coefficient *κ* determines the strength of selection on a change in the cooperative behavior of social partners as a function of how it changes the payoff (i.e., fertility) of the focal individual. If individuals live in a group-structured population, then changes in fertility translate into changes in fitness only after individuals have a chance to disperse and after local competition for resources or breeding patches. This local competition can counteract some (or all) of the positive effect of relatedness on selection for increasing the fertility of a social partner. Thus, low values of scaled relatedness represent either populations with low levels of genetic relatedness or high levels of local competition. High values of scaled relatedness correspond to high levels of genetic relatedness and low levels of local competition. Although determining the value of scaled relatedness that corresponds to a specific demography and genetic relatedness requires calculation (Lehmann and Rousset, 2010; Van Cleve, 2015), one can use scaled relatedness simply as an index of the overall effect of demography and genetic relatedness on selection for social effects on fertility.

**Table 2:** Types of natural selection and the two-player evolutionary games that produce them.

| Selection | Game type | Equilibrium cooperation level | Type of variation |
| --- | --- | --- | --- |
| Directional | Mutualism | Full | No variation |
| Directional | Prisoner's dilemma | None | No variation |
| Diversifying | Stag hunt | Full or none | Among group |
| Balancing | Hawk dove | Mixed / intermediate | Within group |

## 4 The effects of relatedness and synergy on evolutionary games

Using the payoff from Equation (1) in Hamilton’s rule (equation 4), the change in the frequency of the mutant becomes

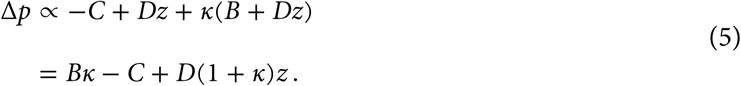

Equation (5) is in effect Hamilton’s rule written in a form that exposes the fertility payoffs of the game and uses scaled relatedness to capture the effect of genetic relatedness and demography on selection for cooperation by social partners. Comparing Equation (5) with the result without relatedness and population structure (equation 2) shows how scaled relatedness *κ* can relax the conditions for the evolution of investment in cooperation. Specifically, Equation (5) shows that the benefit *B* is important only when there is non zero scaled relatedness and that scaled relatedness magnifies the effect of payoff synergy *D*.

One useful way to conceptualize how scaled relatedness changes the conditions under which there is directional, diversifying, or balancing selection on cooperation is to show how scaled relatedness changes the structure of the payoff matrix in Table 1. In deriving the expression for Δ*p* in Equation (2) without scaled relatedness, I used the payoffs from Table 1 to build the payoff function *π* in Equation (1) and then calculated 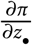. Working backwards from the expression for Δ*p* with scaled relatedness in Equation (5), the payoff matrix that would produce this expression for Δ*p* by simply calculating 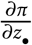 is given in Table 3 (Hamilton, 1971; Grafen, 1979; Hines and Maynard Smith, 1979; Day and Taylor, 1998; Peña et al., 2015). In effect, scaled relatedness shifts the type of evolutionary game by modifying the entries of the payoff matrix. For example, the benefit *B* becomes *B – Cκ* since a focal individual who obtains benefit from a cooperative partner is related to, and hence shares some genetic fate with, their partner where *Cκ* of the partner’s cost translates to the focal individual. Likewise, an individual who cooperates effectively obtains an additional fitness payoff of *Bκ* since individuals it helps are related to it by fraction *κ*. Finally, the payoff of mutual cooperation is increased by 1 + *κ*, which reflects the additional evolutionary success of cooperating with relatives who also cooperate.

**Table 3:** Payoff matrix that accounts for the effect of scaled relatedness *κ*.

|  | C | D |
| --- | --- | --- |
| Payoff to C | $(B - C + D)(1 + \kappa)$ | $-C + B\kappa$ |
| Payoff to D | $B - C\kappa$ | $0$ |

In order to visualize how the combination of payoff synergy and relatedness shifts evolutionary game types, I use Table 3 and Equation (5) in the same way that I used Table 1 and Equation (2) in section 2 to determine what kind of game a particular combination of synergy and relatedness produces. The results of this are shown in the left hand panel (*ρ* = 0.0) of Figure 1 where the fertility cost of cooperation is assumed to be positive (*C* > 0). Synergy is zero (*D* = 0) along the horizontal axis, and the classic result from Hamilton’s rule emerges where directional selection for cooperation (mutualism game) occurs when *Bκ* > *C* and directional selection against cooperation (prisoner’s dilemma) occurs when *Bκ* < *C*. More generally, relatedness greater than *C/D* generates directional selection for cooperation so long as synergy is non-negative. Relatedness *κ* is zero along the vertical axis, and sufficient synergy, *D* > *C*, yields diversifying selection and a stag hunt game. The threshold level of synergy needed to produce diversifying selection decreases with increasing relatedness until *κ* > *C/B*, at which point investment in cooperation is effectively converted from an inclusive fitness cost to an inclusive fitness benefit and selection on investment in cooperation becomes directional and positive. Unlike modifying either relatedness or synergy alone, a combination of negative synergy and relatedness can produce balancing selection (a hawk-dove game). In effect, sufficient positive relatedness (κ > *C/B*) converts the prisoner’s dilemma into a mutualism game, and negative synergy converts the mutualism game into a hawk-dove game. From the perspective of variation in investment (see Table 2), the plot reveals that among group variation in cooperation occurs only when synergy is sufficiently positive and relatedness is not too strong. Within group variation in cooperation occurs only when relatedness is sufficiently strong (*κ* > *C/B*) and synergy is negative.

**Figure 1:**
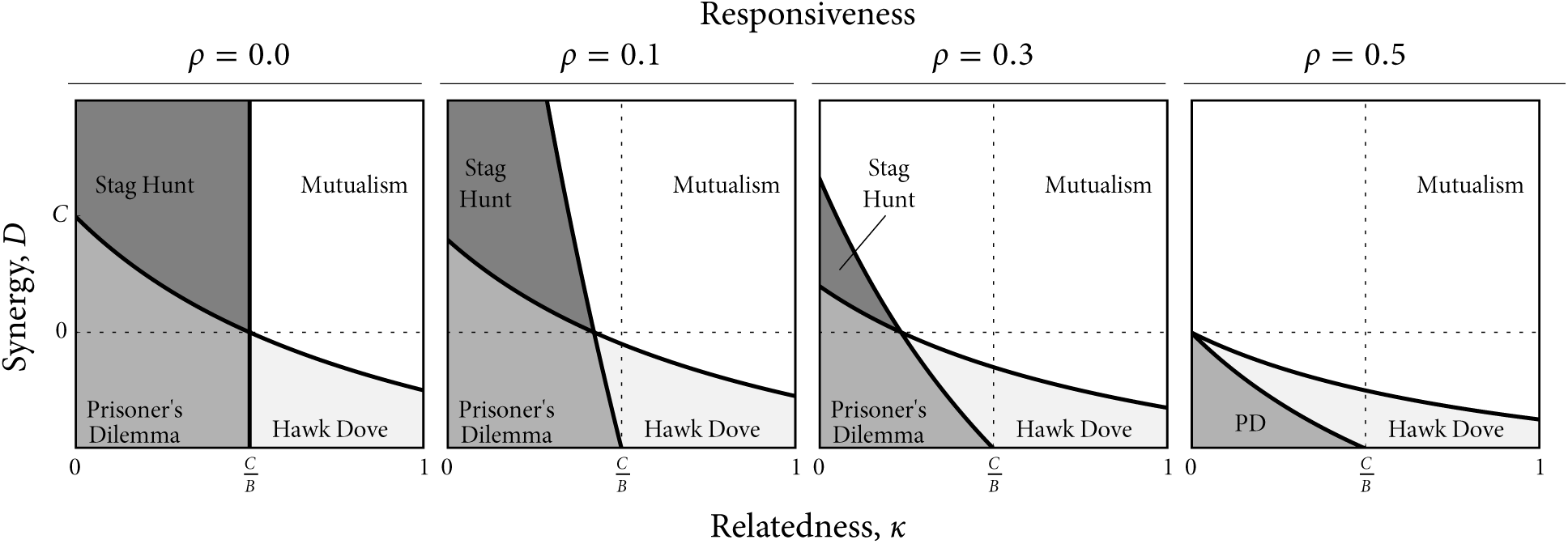
Evolutionary game types produced by the payoff matrix given in Table 5 as a function of synergy *D* and scaled relatedness κ. Responsiveness *ρ* is held fixed in each of the four plots at the value noted. See Table 2 for how these games map to different types of natural selection.

## 5 The effects of responsiveness and synergy on evolutionary games

The payoffs in Tables 1 and 3 (that result in equations 2 and 5) reflect a social interaction where individuals stick to one strategy, cooperate or not. Individuals do not flexibly or plastically respond to the past behavior of their social partners when choosing a strategy; in other words, there is no responsiveness or reciprocity and strategies are fixed by individual genotype. In game theory, behavioral plasticity is studied in the context of repeated or iterated games where individuals choose strategies that determine how way their actions (cooperate or not) depend on the past actions of their social partners. While there is a rich literature that studies which of these repeated strategies yields the best long-term payoff (e.g., Osborne and Rubinstein, 1994; Binmore, 2007; Axelrod and Hamilton, 1981; Boyd and Lorberbaum, 1987; Hilbe et al., 2013; Stewart and Plotkin, 2013), many of those approaches are difficult to integrate with the direct fitness approach used above to incorporate the effect of relatedness. Thus, I will use here a relatively simple model of how individuals can respond to the most recent action of their partner. Analogous to previous work (McNamara et al., 1999; Killingback and Doebeli, 2002; André and Day, 2007; Akçay et al., 2009; Akçay and Van Cleve, 2012; Van Cleve and Akçay, 2014), this model uses a single responsiveness parameter, *ρ*, that measures how likely an individual will respond to its partner by copying their the previous action.

Suppose that individuals have an “intrinsic” action, cooperate or not, that they choose with probability 1 – *ρ* and that they copy the last action of their partner with probability *ρ* (see Supplementary Material A). Individuals choose actions simultaneously and interact repeatedly for long enough so that the mean payoff from the interaction does not depend on the initial rounds (or time steps). Each round, one individual is chosen randomly to update its action and the other individual repeats its action from the previous round. This process of repeated game play can be represented by a transition matrix that specifies the probability that the two individuals use specific actions given their actions in the last round (see equation A.1). In the long run, there is a stationary probability that the two individuals, given their intrinsic strategies, will choose any pair of actions. For each of the four possible pairs of intrinsic strategies that two individuals can choose, the payoff of the interaction can be calculated using the stationary probabilities and the payoffs from Table 1. This leads to the payoff matrix in Table 4 that incorporates responsiveness (see Supplementary Material A).

**Table 4:** Payoff matrix that accounts for the effect of responsiveness *ρ*. See Supplementary Material A for a derivation.

|  | C | D |
| --- | --- | --- |
| Payoff to C | $(B - C + D)(1 + \rho)$ | $-C + B\rho + D\rho$ |
| Payoff to D | $B - C\rho + D\rho$ | $0$ |

Let *z* now measure the probability that an individual’s intrinsic action is cooperation. Using the payoff matrix in Table 4, the change in the frequency of a mutant allele in a population with a resident intrinsic cooperation probability *z* is

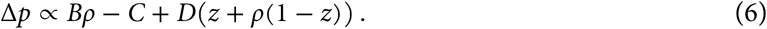

As in the cases with synergy and relatedness, I can use Equation (6) and the payoff matrix in Table 4 to analyze the combined effect of responsiveness and synergy on shifting between evolutionary game types. The leftmost panel in Figure 2 (*κ* = 0.0) shows the results of this analysis. Comparing this panel to the leftmost panel in Figure 1 shows that the combined effects of relatedness and synergy are qualitatively very similar to those of responsiveness and synergy. When responsiveness is zero (*ρ* = 0), enough synergy (*D* > *C*) turns a prisoner’s dilemma with directional selection against cooperation into a stag hunt game with diversifying selection. High levels of responsiveness result in directional selection for cooperation. Intermediate levels of responsiveness create diversifying selection when synergy is intermediate and positive and balancing selection when synergy is intermediate and negative. Unlike relatedness, responsiveness does not have a simple *ρ* > *C/B* threshold for all values of synergy. Rather, this threshold holds only for zero synergy, is relaxed for positive synergy (mutualism game possible for *ρ* < *C/B*) and is stricter for negative synergy (prisoner’s dilemma game possible for *ρ* > *C/B*). Nevertheless, the interpretation is similar in that increasing responsiveness converts the direct fitness cost of cooperation into a direct fitness benefit, which results in directional selection for cooperation when synergy is positive and first diversifying selection and then directional selection for cooperation when synergy is negative. Focusing on when there is variation in cooperation, responsiveness generates between population variation via diversifying selection when synergy is sufficiently positive and responsiveness is low and within group variation via balancing selection when synergy is moderately negative and responsiveness is high. Compared to relatedness, responsiveness has a smaller scope for variation in cooperation via either diversifying or balancing selection.

**Figure 2:**
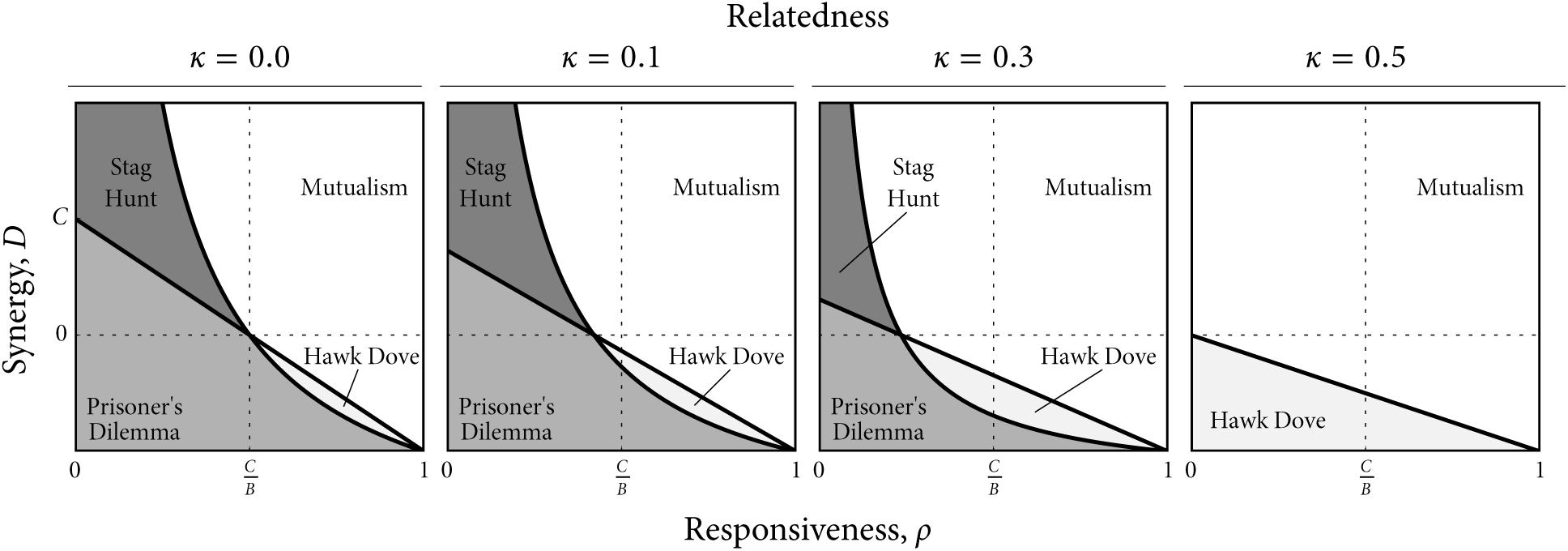
Same as Figure 1 except results are plotted as function of responsiveness *ρ* and synergy *D* with scaled relatedness *κ* held constant in each plot at the value noted.

## 6 The combined effects of relatedness, responsiveness, and synergy

Equation (6) and Table 4 correspond to a social interaction in an unstructured population of unrelated individuals who are responsive to their social partners with probability *ρ*. Additionally accounting for relatedness involves using the payoffs from Table 4 in Equation (4), which yields

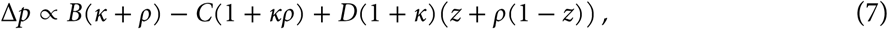
which is equivalent to previous results without synergy (*D* = 0) (Akçay and Van Cleve, 2012; Van Cleve and Akçay, 2014). As in section 4, Equation (7) can be generated by calculating only 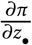 using an augmented payoff matrix, which is given in Table 5. Using the payoff matrix in Table 5 and Equation (7), I show the joint effect of relatedness, responsiveness, and synergy in the three rightmost panels in Figures 1 and 2. Both figures show that increasing relatedness and responsiveness together can easily lead to positive directional selection for cooperation, particularly if synergy is zero or positive. Since diversifying selection relies on cooperation being individually costly, a large enough combination or relatedness and responsiveness eliminates the possibility of diversifying selection by effectively making cooperation individually beneficial. In contrast, negative synergy can still generate balancing selection under these conditions so long as neither relatedness nor responsiveness is too close to one. This suggests that variation in investment in cooperation is most likely to be within groups when relatedness or responsiveness is significantly greater than zero.

**Table 5:** Payoff matrix that accounts for the effects of both scaled relatedness and responsiveness.

|  | C | D |
| --- | --- | --- |
| Payoff to C | $(B - C + D)(1 + \kappa)(1 + \rho)$ | $B(\kappa + \rho) - C(1 + \kappa\rho) + D\rho(1 + \kappa)$ |
| Payoff to D | $B(1 + \kappa\rho) - C(\rho + \kappa) + D\rho(1 + \kappa)$ | $0$ |

## 7 The effect of social group size and relatedness

Extending the model from pairwise social interactions to social interactions involving larger groups adds substantial complexity since it expands the payoff matrix to account for all possible group compositions of who cooperates and who does not. This expanded payoff matrix is given in Table 6 for a group of size *n* where *u_k_* and *v_k_* are the payoffs an individual receives for cooperating and not cooperating, respectively, when *k* other individuals in their group cooperate and *n–k*–1 other individuals do not cooperate. There are many ways that payoff can change as the number of cooperators in the social groups changes and this leads to the possibility of many different stable and unstable group compositions (i.e., fractions of cooperators or values of the mixed strategy 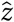; Motro, 1991; Bach et al., 2006; Gokhale and Traulsen, 2010; Peña et al., 2014, 2015). In other words, *n*-player games generically have a more complex set of outcomes than two-player games, and thus, unlike two-player games, cannot be easily related uniquely to directional, diversifying, and balancing selection as described in section 2. In order to determine the effect of increasing social group size without this additional complexity, I borrow from Peña et al. (2015) who demonstrated an elegant description of *n*-player payoffs in cooperation (or “public goods”) games that yield only the four outcomes obtained in two-player games (see Table 2). The crux of the approach is that if increasing the number of individuals in the group who cooperate always increases the benefit available to the group as a public good, then the only outcomes possible are the ones obtained in two-player games (Peña et al., 2015). This approach then allows one to study the effect of increasing group size as well as relatedness, responsiveness, and synergy. I present results for relatedness and synergy below and leave responsiveness for future work.

**Table 6:** Payoff matrix for a social interaction involving *n* individuals. Fixed total Fixed per capita

| Partners playing C | $n - 1$ | … | 1 | 0 |
| --- | --- | --- | --- | --- |
| Payoff to C | $u_{n-1}$ | … | $u_1$ | $u_0$ |
| Payoff to D | $v_{n-1}$ | … | $v_1$ | $v_0$ |

In the *n*-player case, *C* is still the individual cost of cooperation. In order to control for the potential that total group benefit changes as group size increases, the payoff for cooperation when all social partners (*n*–1 of them) also cooperate is set at *B–C*+*D*, which is same as for the two-player game. Further, assume that the total additive benefit when all individuals cooperate is *B*, which means that each investment by a social partner produces *B/*(*n* – 1) additive benefit. For a fixed group size, the public good or total benefit available to the group increases with each additional individual who cooperates. Assume that the *k*^th^ investment in cooperation contributes the additive benefit, *B*/(*n* – 1), times a synergy parameter *λ*^*k*–1^. Thus, two individuals who cooperate produce a total benefit of *B*/(*n* – 1)(1 + *λ*), three individuals produce *B*/(*n* – 1)(1 + *λ* + *λ*^2^), and so on (Peña et al., 2015, equation 12). When *λ* = 1, there is no synergy and individual investments in cooperation contribute additively to the total benefit, which corresponds to *D* = 0 in the two-player game. When *λ* > 1 and *λ* < 1, there is positive synergy (*D* > 0) and negative synergy (*D* < 0), respectively. The “Fixed total additive benefit” column in Table 7 shows the payoffs for this scenario and the relationship between the two-player and *n*-player synergy parameters *D* and *λ*.

**Table 7:** Two different n-player payoff scenarios for the payoff matrix in Table 6 and the corresponding relationship between the *n*-player and two-player synergy parameters. In the first scenario, the total additive benefit in the social group is fixed at *B* and the per capita benefit is *B/*(*n –* 1). In terms of the parameters of Peña et al. (2015, Table 1), *β* = *B/*(*n –* 1) and *γ* = *C* + *B/*(*n –* 1). The additive per capita (per individual) benefit is fixed at *B* in the second scenario leading to a total additive benefit (*n –* 1)B, *β* = *B* and *γ* = *B* + *C*.

|  | Additive benefit type |  |
| --- | --- | --- |
|  | Fixed total | Fixed per capita |
| Payoff to C ($u_k$) | $-C + \frac{B\lambda}{n-1} \left( \frac{\lambda^k - 1}{\lambda - 1} \right)$ | $-C + B\lambda \left( \frac{\lambda^k - 1}{\lambda - 1} \right)$ |
| Payoff to D ($v_k$) | $\frac{B}{n-1} \left( \frac{\lambda^k - 1}{\lambda - 1} \right)$ | $B \left( \frac{\lambda^k - 1}{\lambda - 1} \right)$ |
| Additive synergy $D$ | $\frac{B}{n-1} \left( \frac{\lambda^n - 1}{\lambda - 1} - n \right)$ | $B \left( \frac{\lambda^n - 1}{\lambda - 1} - n \right)$ |

The results of Peña et al. (2015) can be used to determine which of the four types of selection on cooperation and their corresponding game types is produced by a specific combination of *n, B, C*, and *D*. Using these results (Peña et al., 2015, Table 2), I plot in Figure 3 the type of selection and evolutionary game as a function of relatedness, synergy, and social group size. Figure 3 shows that increasing social group size given this payoff scenario has a remarkably small effect on how relatedness and synergy determine the type of selection on cooperation. The most significant effect of increasing group sizes is when relatedness is close to zero: larger group sizes require higher levels of synergy to switch directional selection against cooperation to diversifying selection. Otherwise, increasing relatedness retains its strong effect on shifting the inclusive fitness cost of cooperation to an inclusive fitness benefit when *κ* > *C/B*, which generates directional selection for cooperation when synergy is non-negative and balancing selection when synergy is sufficiently negative.

**Figure 3:**
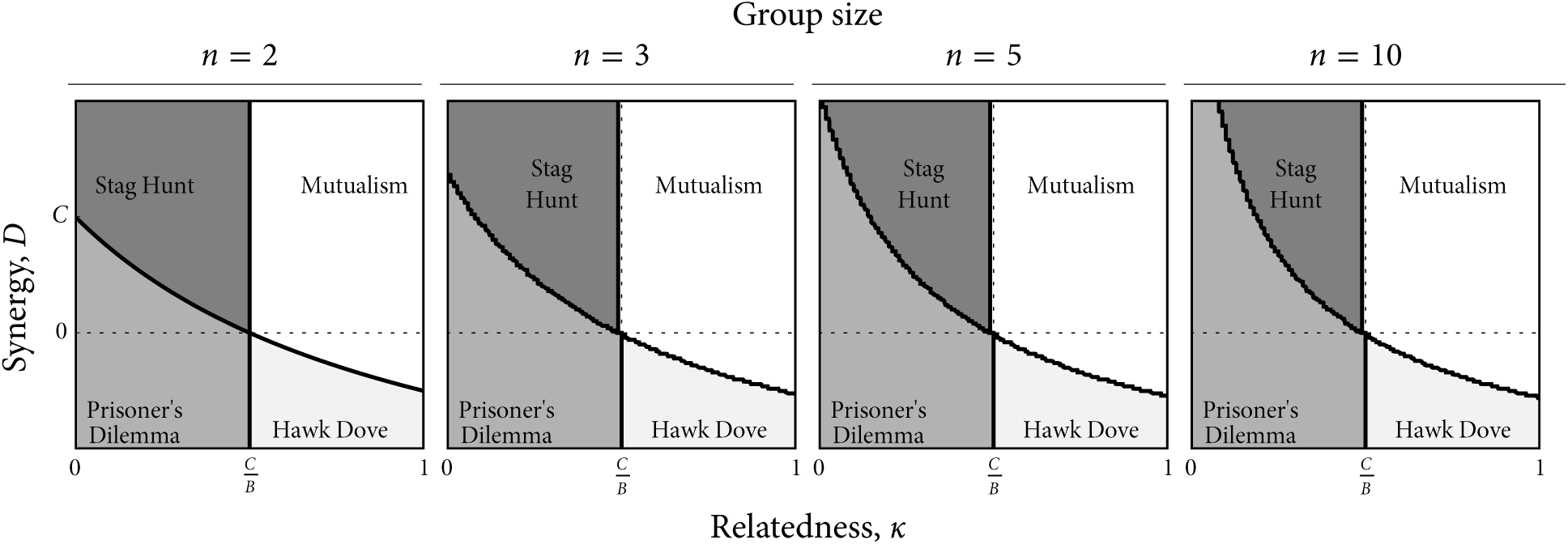
Evolutionary game types as a function of scaled relatedness and synergy where each plot is a different social group size. The payoffs used to generate these plots are in Table 7 under the heading “Fixed total additive benefit”.

Fixing the total additive benefit at *B* and allowing the per capita benefit to decrease with group size (*B*/(*n*–1)) in Figure 3 appears to produce an *n*-player game whose evolutionary outcomes are remarkably similar to the two-player case. This similarity evaporates if these previous assumptions are changed. For example, I can fix the per capita benefit at *B*, which means it does not change with group size, and allow the total additive benefit, (*n* –1)*B*, to increase with group size. The payoffs for this case are given in the “Fixed per capita additive benefit” column in Table 7. Notably, fixing the per capita benefit at *B* is equivalent to setting the cost to *C*/(*n* – 1) and shrinking it with group size since what matters for the group-structured demography used here is relative, not absolute, payoff. Using these payoffs, Figure 4 shows the type of selection on investment in cooperation as a function of relatedness and synergy for different social group sizes. Allowing the total group benefit to increase with group size strongly increases the effect of relatedness; the threshold relatedness required to shift investment in cooperation from an inclusive fitness cost to an inclusive fitness benefit decreases as group size increases. Thus, much weaker levels of relatedness can generate directional selection for cooperation when synergy is positive and balancing selection when synergy is sufficiently negative. The scope for directional selection against cooperation and diversifying selection is reduced rapidly as group size increases since the threshold relatedness decreases rapidly. As group size increases, synergy must be increasingly negative in order to generate balancing selection for a given value of relatedness. Though this decreases the scope for balancing selection, it remains much more likely than diversifying selection as group size increases. Thus, variation in investment cooperation, if it exists in this particular scenario, is more likely within groups as group size increases.

**Figure 4:**
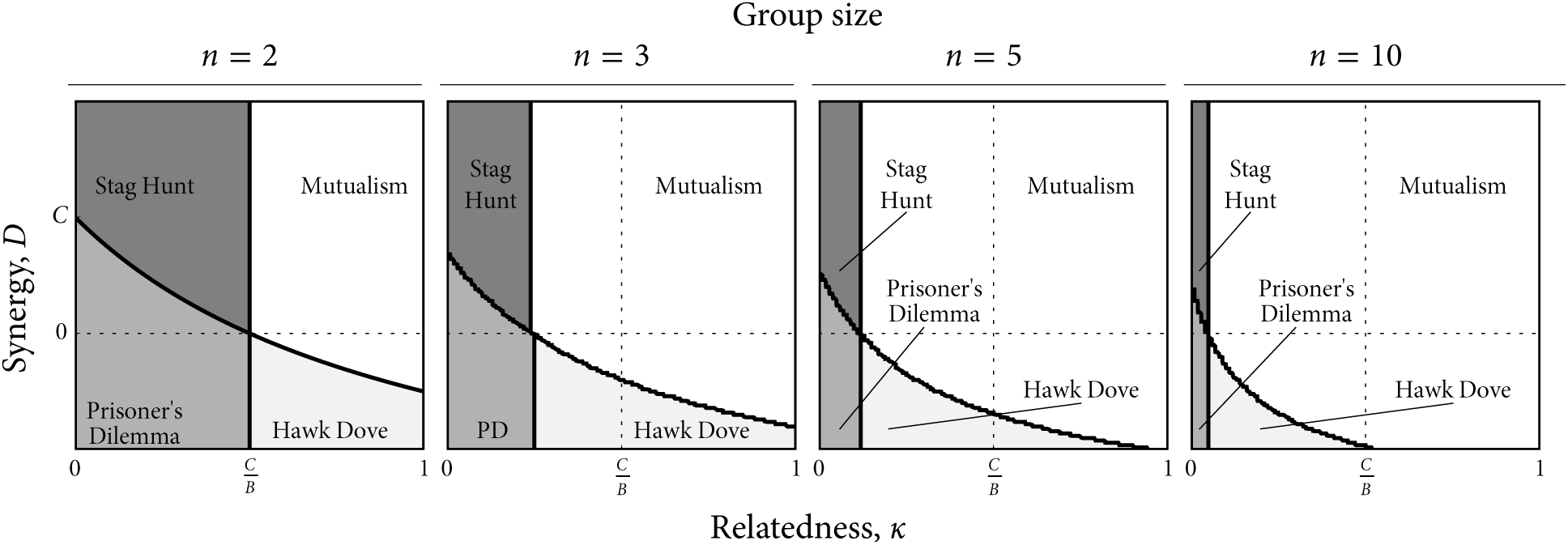
Same as Figure 3 except the payoffs used to generate the plots are in the “Fixed per capita additive benefit” column in Table 7. Note that before plotting, the payoffs have all been normalized by *n –* 1, which keeps the total additive payoff and the synergy *D* on the same scale as in Figure 3.

## 8 Discussion

Understanding the combined role of relatedness, responsiveness, and synergy in promoting the evolution of cooperation has been hindered by a lack on integration of these mechanisms in a common theoretical framework. Here, I show how kin selection theory and evolutionary game theory provide an approach that reveals how the effects of relatedness, responsiveness, and synergy can be characterized in terms of the four basic two-player games and how these games create directional, balancing, or diversifying selection on cooperation (Table 2). The framework here shows how relatedness, responsiveness, and synergy created different types of selection on cooperation by effectively transforming the structure of the payoff matrix (e.g., Table 5). Past work has also emphasized that demographic, behavioral, and ecological parameters can alter the structure of the payoff matrix (Taylor and Nowak, 2007; Van Cleve and Akçay, 2014; Taylor and Maciejewski, 2012), and that such parameters may themselves evolve (Akçay and Roughgarden, 2011; Stewart and Plotkin, 2014).

While the importance of synergy in shaping variation in cooperation has been highlighted in the past (e.g., Fletcher and Doebeli, 2006; Hauert et al., 2006; Cornforth et al., 2012), my results here emphasize the special role that synergy has vis-à-vis other factors that are conducive to investment in cooperation like relatedness, responsiveness, and group size. Specifically, responsiveness and relatedness are primarily important in creating conditions generally conducive to investment in cooperation whereas synergy determines whether there is variation or not. Synergy creates frequency-dependent selection where frequency in the framework here refers to the probability of investing in cooperation *z*. With additive benefits and costs of investing in cooperation and no synergy, selection is frequency independent and directional and leads to either full cooperation or no cooperation. Positive synergy generates positive frequency-dependent selection, which makes the full cooperation equilibrium (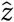 = 1) stable when synergy is strong enough. If the no cooperation equilibrium (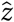 = 0) is also stable because relatedness and responsiveness are low and cooperation is a fitness cost, then positive synergy can create diversifying selection and a stag-hunt game. If the no cooperation equilibrium is unstable because relatedness or responsiveness is high enough (cooperation is a fitness benefit), then positive synergy enhances convergence to the full cooperation equilibrium. In contrast, negative synergy creates negative-frequency dependent selection, which if negative enough leads to instability of the full cooperation equilibrium even when relatedness or responsiveness is high. High relatedness or responsiveness makes the no cooperation equilibrium unstable, which leaves a mixed-strategy equilibrium (0 < 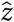 < 1) stable due to balancing selection.

Integrating relatedness, responsiveness, and synergy in a single framework also reveals new evolutionary pathways between closely related species that exhibit variation in investment in cooperation. Variation in cooperation between groups, populations, or species can be created by diversifying selection due to a stag-hunt game whereas within group variation can be created by balancing selection due to a hawk-dove game. These two game types are often seen as the result of disparate ecological scenarios since a hawk-dove game in simple scenarios without relatedness and responsiveness requires that cooperation yields an individual payoff benefit (*C* < 0) whereas a stag-hunt requires that cooperation exacts an individual cost (*C* > 0). Explaining how closely related species evolved both within and between group variation in cooperation then requires explaining how this one fundamental payoff parameter could have changed. By combining the effects of relatedness, responsiveness, and synergy, I show how both within and between group variation in cooperation are both possible even when cooperation has an individual payoff cost. Low levels of relatedness and responsiveness translate the individual payoff cost into a fitness cost and between group variation in cooperation is then possible when synergy is sufficiently positive. High levels of relatedness and responsiveness create a fitness benefit despite the payoff cost, which allows for within group variation in cooperation when synergy is sufficiently negative. Thus, it is possible to create conditions for within or between group variation in investment in cooperation by altering the economics of generating benefits (i.e., synergy) and either the population demography (e.g., levels of dispersal that shape relatedness) or the amount of behavioral plasticity (e.g., behavioral responsiveness).

Using this framework, I also show how changes in social group size are alone not guaranteed to have much effect on how relatedness, responsiveness, and synergy affect investment in cooperation. When the total additive benefit of cooperation is fixed (the per capita additive benefit decreases with group size), increasing group size has remarkably little effect on the conditions necessary for selection to favor cooperation or variation in cooperation. The only notable effect is that low levels of relatedness require higher levels of positive synergy to generate diversifying selection. When the per capita additive benefit of cooperation is fixed and the total additive benefit increases with group size, larger groups are much more conducive to investment in cooperation. In particular, the threshold level of relatedness necessary for cooperation to be a fitness benefit decreases quickly as group size increases. Increasing group size in this scenario also decreases the scope for diversifying selection, which leaves balancing selection and within group variation the more likely outcome if cooperation levels vary.

In addition to providing conceptual insights, this framework can also be tested empirically by measuring levels of relatedness, responsiveness, and synergy. Measuring levels of genetic relatedness is routine (Rousset, 2002) and there are sophisticated statistical methods based on the animal model of quantitative genetics that can properly account for identity by descent as measured by pedigrees and the effect of that identity on traits such as cooperation (Lynch and Walsh, 1998; Wilson et al., 2010). Further developments of the animal model called “indirect genetic effects” (IGE) also allow estimation of responsiveness coefficients by measuring an additional regression coefficient (Moore et al., 1997; McGlothlin et al., 2010; Akçay and Van Cleve, 2012; McGlothlin et al., 2014). There has been less emphasis on measuring the degree of nonadditivity in the production of benefits from the social interaction (i.e., synergy). This can likely be accomplished by adding nonlinear terms to IGE regression models. Manipulative experiments can also measure synergy by changing the investment level of individuals or the number of individuals investing at a certain level and measuring the resulting benefits obtained by the social group. Thus, future empirical studies should be able to simultaneously measure relatedness, responsiveness, and synergy and test predictions regarding the evolved levels of cooperation and variation in levels of cooperation. Such predictions are particularly important for understanding conditions that favor or disfavor the evolution behavioral syndromes in animal populations (Sih et al., 2004; Wolf and Weissing, 2012; Jandt et al., 2014).

## 9. Acknowledgements

I would like to thank the following funding sources: USA National Science Foundation IOS-1634027, Society of Integrative and Comparative Biology Division of Animal Behavior, Division of Comparative Endocrinology, Division of Ecology and Evolution, and Division of Neurobiology, Neuroethology, and Sensory Biology. I would also like to thank the organizers of the symposium, Ben Dantzer and Dustin Rubenstein.

## Supplementary Material A: Two-player responsiveness model

Suppose that individuals engage in pairwise interactions where they simultaneously choose either to cooperate (C) or to not cooperate (D) and obtain payoffs given in Table 1. At the beginning of every interaction, each individual is either (i) an intrinsic cooperator who cooperates by default with probability 1–*ρ_D_* or responds to its partner by copying the partner’s last action with probability *ρ_C_* or (ii) an intrinsic non-cooperator who does not cooperate by default with probability 1 – *ρ_D_* and copies its partner’s last action with probability *ρ_D_*. Since my analysis is focuses on understanding how responsiveness *ρ* affects selection on investment in cooperation and not on understanding how *ρ* evolves intrinsically, I set *ρ_C_* = *ρ_D_* = *ρ*. Since individuals only require knowledge of their partner’s last action, the model is Markovian and the state space of the model consists only of the four possible current states, (D,D), (D,C), (C,D), and (C,C), where the elements are the action of the focal individual and its partner, respectively. Further assume that each time step only one of the two individuals has an opportunity to update its action (i.e., choose its “intrinsic” action with probability 1 *– ρ* and copy its partner with probability *ρ*). This is kind Markov model is called a “quasi birth-death process” (Latouche and Ramaswami, 1999) and is simpler than alternative formulations when the model is extended from pairwise to n-player interactions (Van Cleve, unpublished).

Since individuals can act as either intrinsic cooperators or non-cooperators, there are four possible pairwise interactions and each interaction has a transition matrix describing the Markov chain. Each Markov chain is irreducible and aperiodic (for 0 < *ρ* < 1) and thus has a unique stationary distribution that describes the probability of each pairwise state in the long run for the corresponding social interaction. Independent of the value of ρ, an interaction between two intrinsic cooperators has a stationary distribution where both individuals always cooperate; likewise, two intrinsic defectors both always defect in the long run. The transition matrix for an interaction between a focal intrinsic cooperator and a non-cooperator social partner is

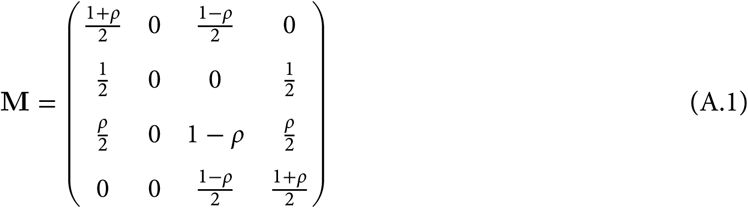
where each element specifies the probability that the row state leads to the column state and the order of states in the rows and columns is (D,D), (D,C), (C,D), and (C,C). The stationary distribution *ν* of M is

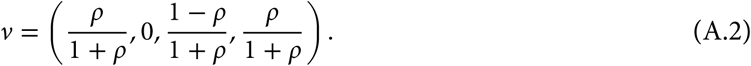
When *ρ* = 0, the stationary distribution *ν* becomes (0,0,1,0), which means that the intrinsic cooperator always cooperates and the intrinsic non-cooperator never cooperates and corresponds to the model without responsiveness or reciprocity. When *ρ* = 1, the stationary distribution is *ν* = (*1/2*,*0*,*0*,1/2), which reflects perfect responsiveness where individuals who both begin cooperating or not cooperating always continue cooperating or not cooperating. Pairs that begin with one individual cooperating and the other not cooperating end up either both cooperating or not cooperating with equal probability. The transition matrix for a focal intrinsic non-cooperator interacting with an intrinsic cooperating partner is the transpose of M from equation (A.1) and the stationary distribution *ν* is also given by equation (A.2) except with the labels of the states inverted (C replaced by D and vice versa).

For simplicity, I assume that payoffs are accrued only at the stationary distribution, which means that the interaction is long enough so that transient payoffs at the beginning of the game can be discarded. Payoffs for each of the four pairwise combinations of intrinsic cooperators and non-cooperators can then be calculated by using the stationary distributions of the four transition matrices and multiplying each element of distribution with the corresponding payoff from Table 1. This results in Table A.1, which simplifies to Table 1 when *ρ* = 0. Since Nash equilibria are invariant under linear transformations of payoffs (Straffin, 1993, p. 50), the entries of Table A.1 can be multiplied by 1 + *ρ*, which generates Table 4.

**Table A.1:** Payoff matrix for the two-player responsiveness model.

|  | intrinsic C | intrinsic D |
| --- | --- | --- |
| Payoff to intrinsic C | $(B - C + D)$ | $\frac{-C+B\rho+D\rho}{1+\rho}$ |
| Payoff to intrinsic D | $\frac{B-C\rho+D\rho}{1+\rho}$ | 0 |

